# Biological characteristics of two mesenchymal stem cell lines isolated from the umbilical cord and adipose tissue of a neonatal common hippo (*Hippopotamus amphibius*)

**DOI:** 10.1101/2021.12.26.474212

**Authors:** Jinpu Wei, Xiuxiu Dong, Bo Wang, Yajiang Wu, Wu Chen, Zhijun Hou, Chen Wang, Tao Wang

**Author notes:** **Correspondence Name:** Tao Wang; **E-mail:**, **Name:** Chen Wang; **E-mail:**.

## Abstract

Mesenchymal stem cells (MSCs) are multipotent adult stem cells and can be isolated from many tissues of the body. Due to their potentials to treat various diseases and be applied in animal breeding, MSCs have been isolated and identified regarding their biological properties. Common hippos (*Hippopotamus amphibius*) are a vulnerable species and yet the cryopreservation of their genetic materials is scare. In this study, we successfully established two MSC lines (UC-MSCs and AT-MSCs) from the umbilical cord and adipose tissue of a neonatal common hippo and comparatively described their features. Both UC-MSCs and AT-MSCs showed fibroblastoid morphology and could be continuously passaged for over 17 passages without dramatic signs of senescence. The cell cultures had normal chromosome composition, say, 17 pairs of autosomes and 1 pair of X chromosomes. UC-MSCs and AT-MSCs displayed similar gene expression profiles. They were positive for CD45, CD73, CD90 and CD105 and negative for HLA-DR. They demonstrated stemness maintenance by expression of classical stem cell markers. UC-MSCs and AT-MSCs manifested different differentiation potentials into other cell lineages. In summary, these two cell lines demonstrated the essential properties of mesenchymal stem cells and might play a role in the future research.

## 1. Introduction

Mesenchymal stem cells (MSCs) are a heterogeneous cell population that could be isolated from many tissues of the body, e.g., bone marrow, umbilical cord, placenta, adipose tissue, dental pulp and even circulating blood (Hendijani, 2017; Iser et al., 2014; Sato et al., 2016; Wei et al., 2016). It is acknowledged that MSCs have the following common characteristics. They exhibit a feature of adherence to the culture plates and are morphologically similar to fibroblasts. And they generally express certain cell surface proteins, e.g., CD73, CD90 and CD105, and lack the expression of hematopoietic and endothelial cell markers, e.g., CD34, CD45 and HLA-DR. Besides, MSCs maintain the potentials of self-renewal and multi-differentiation into tissue-specific lineages.

So far, there have been substantial research papers reporting the isolation and characterization of MSCs from different tissues of various animal species (e.g., Bourzac et al., 2010; Cortez et al., 2018; Lyahyai et al., 2012). What is of interest is that MSCs behaved somewhat inconsistently across different studies. To be detailed, the acquired MSCs might vary with regard to e.g., the proliferative capacity, surface marker expression and the differentiation potentials (Carrade and Borjesson, 2013; Iser et al., 2014; Uder et al., 2018). The observed variations are likely to be a combinatorial result from species, sources of MSCs, isolation methods, propagation techniques, antibody availability and etc. In fact, many comparative projects were conducted to shed light on how the phenotypical and functional properties could be affected by the abovementioned factors (Bourzac et al., 2010; Clark et al., 2016; Reich et al., 2012; Sun et al., 2020).

MSCs have attracted considerable attention within the field of cell-based therapies due in part to their capability of differentiation and in part to their secretory properties to treat various diseases (Al-Ghadban and Bunnell, 2020; Galland and Stamenkovic, 2020; Lelek and Zuba-Surma, 2020; Tsiapalis and O’Driscoll, 2020). In addition, existent evidence demonstrates great values of MSCs for animal assisted reproduction. Addition of extrinsic factors such as BMP4, TGFβ1 and retinoic acid into the cultured MSCs could lead to the induction of germ cells *in vitro* (Afsartala et al., 2016; Cortez et al., 2018; Ghasemzadeh-Hasankolaei et al., 2014; Wei et al., 2016). MSCs could also be applied as nuclear donors for the somatic nuclear transfer and successful attempts have been achieved for pigs and cattle (Colleoni et al., 2005; Li et al., 2013; Yang et al., 2016).

The common hippopotamus or hippo (*Hippopotamus amphibius*) is well-known for its bulky body and semi-aquatic habit. Despite their aggressive nature, the common hippo population in the wild declines rapidly in the recent years (Lewison, R. & Pluháček, 2017). As a result, the common hippo is described as vulnerable in the IUCN Red List of Threatened species and included in the Appendix II of CITES (Lewison, R. & Pluháček, 2017). In this regard, preserving the genetic resources to maintain the diversity of the genetic pool is necessary for their potential applications such as animal breeding and cloning. To the best of our knowledge, there has been no publication on isolating and cryopreserving MSCs from the common hippo so far.

In this project, we tried to culture and characterize two mesenchymal stem cell lines derived from a neonatal common hippo. To this end, we obtained the umbilical cord and subcutaneous adipose tissue from the dead calf for MSCs culture *in vitro* and identified the biological characteristics of the isolated cells using a variety of testing techniques.

## 2. Material and methods

### 2.1. Tissue collection from a common hippo

This study was approved by the Institutional Review Board on Bioethics and Biosafety of China National GeneBank (No. CNGB-IRB 2001-T1) and the procedures involving sample collection were in compliance with applicable regulations and ethical principles. The umbilical cord from a neonatal common hippo was collected and the adipose tissue beneath the skin was taken later that day when the newborn calf died of unknown cause. The sampled tissues were maintained in the preserving solution consisting of DMEM (Invitrogen, 12430054), 5×penicillin/streptomycin (P/S) (Gibco, 15140122) and 2×amphotericin B (Sigma, A2942) at 4°C.

### 2.2. Isolation and culture of MSCs

Classical explant culture and enzymatic digestion were adopted to isolate MSCs from the umbilical cord and subcutaneous adipose tissue, respectively. To be detailed, the umbilical cord and adipose tissue were initially incubated with saline solution added with 10×P/S and 2×amphotericin B for 10 min. After dissected into small explants measuring 1 mm2 in size, they were rinsed with sterile saline solution. The umbilical cord explants were subsequently seeded into the culture flasks and supplemented with complete culture medium composed of DMEM/F12 (Gibco, C11330500BT), 20% fetal bovine serum (FBS) (Biowest, S181F), 1×P/S and 1×amphotericin B for primary culture. On the other hand, the adipose tissue explants were digested with collagenase IV (Invitrogen, 17104-019) for 30 min at 37°C to obtain the cell suspension which was then filtered via 100-μm cell strainers and centrifuged. The cell pellets were resuspended in fresh complete culture medium and transferred to culture flasks.

The isolated cells were cultured at 37°C in a humidified CO2 incubator. When the cell confluence reached ~90%, the cultures were treated with 0.25% trypsin solution (Thermo, 15050057) at 37°C for 2-3 min until the cells detached from the bottom of the culture flasks. The cell suspension was collected, centrifuged and resuspended in new complete culture medium. Subculture was performed with the seeding density of 1×10^4^ viable cells/cm2 in new culture flasks.

Potential microorganism contamination was constantly detected via direct observations of bacterial/fungi growth under a light microscope. Besides, BacT/ALERT 3D Microbial Identification System and PCR Mycoplasma Detection Set (Takara, 6601) were applied to detect bacterial/fungi and mycoplasma contamination, respectively.

### 2.3. Cryopreservation and recovery

Cryopreserving the MSCs of early passages was conducted when the cell confluence was of ~90%. The cells were collected after incubation with trypsin solution and resuspended in freezing medium made up of 90% FBS and 10% DMSO (Sigma, D8418). The cell viability was measured based on trypan blue staining method on an automated cell counter. The final concentration of cell suspension in cryogenic vials was 2×10^6^ viable cells/mL.

After a short-term cryopreservation in liquid nitrogen, the frozen cells were rapidly warmed at 37°C in a water bath. The cells were then centrifuged and resuspended in fresh complete culture medium. After the viability was detected, the cells were re-seeded into the culture flasks for serial passaging.

### 2.4. Growth curve analysis

MSCs growth dynamics were assessed according to CCK-8 method that is a convenient colorimetric assay to determine the number of viable cells. In brief, a total of 1.5×10^3^ viable cells per well were initially inoculated in the 96-well plate and six replicates were adopted. The absorbance values at the wavelength of 450 nm were detected under a microplate reader after the cells were treated with CCK-8 solution (Beyotime, C0038) for 1 h. A growth curve was plotted based on 7-day continuous measurements, with culture time on the X-axis and absorbance value on the Y-axis. Each point was plotted as Mean ± SD using Microsoft Excel.

### 2.5. Karyotyping

Cryopreserved MSCs were thawed and re-seeded in the 6-well culture plate for chromosome analysis. When the cells were in a rapid-proliferation state, the colcemid solution (Santa Cruz, sc-202550) was added with the final concentration of 1 μg/mL and incubated with the cultures at 37°C for 6 h. Incubation of the cell cultures with pre-warmed hypotonic solution (0.075 M KCl) was then performed at 37°C for 10 min. After that, the cells were fixed with cold fixative solution which was composed of methanol and glacial acetic acid (1:3) three times. The obtained cell suspension was thereafter dropped onto cold clean glass slides and stained with Giemsa solution (Solarbio, G1010) after drying at 85°C for 4 h. The chromosomes were imaged under a light microscope and ~30 spreads for each cell culture were analyzed.

### 2.6. Immunofluorescence staining of cell surface markers and intracellular proteins

Cells cultured on the coverslips in 24-well plates were fixed with 4% paraformaldehyde for 15 min, followed by washing with phosphate-buffered saline (PBS) three times. The cells were thereafter incubated with 5% goat serum (Boster, AR0009) (for cell surface marker detection) or incubated with 5% goat serum added with 0.2% TritonX-100 (for intracellular protein detection) for 1 h and rinsed with PBS. Incubation with primary antibodies against cell surface markers (CD45, CD73, CD90, CD105 and HLA-DR) (Abcam, ab30470; ab175396; ab225; ab230925; ab136320) or intracellular markers (C-MYC, KLF-4, NANOG, OCT-4 and SOX2) (Santa Cruz, Sc-40; Abcam, ab214666; ab106465; ab18976; ab97959) was performed in the dark at 4°C overnight. The antibodies were diluted according to the user manual. Cells in the blank control group were incubated with PBS only. After rinsed, the cells were incubated with appropriate secondary antibodies in the dark for 4 h. Following counterstaining with DAPI (Beyotime, C1006) for 8 min, the cells were washed and mounted with anti-quench mounting medium for imaging. Unless otherwise noted, the experiments were conducted at room temperature.

### 2.7. Trilineage differentiation

The abilities of the isolated MSCs to differentiate into chondrocytes, osteoblasts and adipocytes were determined, respectively. Commercial trilineage differentiation kits (Chembio, CHEM-200028; CHEM-200029; CHEM-200030) were applied and the differentiation of MSCs was achieved under the instructions of the user manual. For chondrogenesis, MSCs were centrifuged gently and cultured in 15-mL centrifugation tubes added with chondrogenic differentiation medium. As required, the differentiation medium was changed every second day. After 27-day induction, the cell spheres were fixed, dehydrated and embedded in paraffin for subsequent staining with Alcian Blue solution. For osteogenic differentiation, MSCs were inoculated in the gelatin-coated 6-well plates with a density of 4×10^4^ viable cells/cm^2^. Osteogenic differentiation medium was supplemented when the cells reached a confluence of 60% and exchanged every 3 days. 5 weeks later, the cell cultures were fixed and stained with Alizarin Red to show the appearance of osteoblasts. For adipogenic induction, MSCs were seeded into the gelatin-coated 6-well plates with a cell concentration of 3×10^4^ cells/cm^2^. When the cell confluence reached ~90%, adipogenic differentiation solution A was added and then replaced by differentiation solution B after 72 h. 24 h later, the culture medium went back to differentiation solution A. Repeat these two procedures until apparent lipid droplets could be observed. Finally, the cell cultures were fixed and incubated with Oil Red O solution.

## 3. Results

### 3.1. Isolation and culture of MSCs from the common hippo

We acquired the umbilical cord and subcutaneous adipose tissue of a neonatal common hippo and took advantage of the explant culture and enzymatic digestion methods to establish two cell lines. The cell cultures were designated as Umbilical Cord-Derived Mesenchymal Stem Cells (UC-MSCs) and Adipose Tissue-Derived Mesenchymal Stem Cells (AT-MSCs), respectively. The detailed information about the cell lines has been deposited on the online Biological Resource Center of Plants, Animals and Microorganisms of China National GeneBank Database (CNGBdb) (https://db.cngb.org/brc/) and thus could be accessed via the catalogue number, CNGBCCAC00085 and CNGBCCAC00087.

After 7 days of primary culture, a plenty of long and thin cells emerged surrounding the umbilical cord tissue explants. The cells exhibited a polygonal or spindle-like shape with a few processes reminiscent of fibroblast cells (Figure 1A). Similarly, the obtained cells from the digested adipose tissue also showed a fibroblastoid morphology and adherent property (Figure 1D). Around 2 weeks after the initial seeding, UC-MSCs reached a confluence of ~90%. Comparatively, AT-MSCs reached the confluence after 10 days of primary inoculation. Both cell cultures proliferated rapidly and had a viability of over 92% when they were collected for cryopreservation. After the freezing-and-thawing process, UC-MSCs and AT-MSCs still maintained a healthy state. They presented a normal morphological feature and had a high cell viability, say, over 90%.

**Figure 1.**
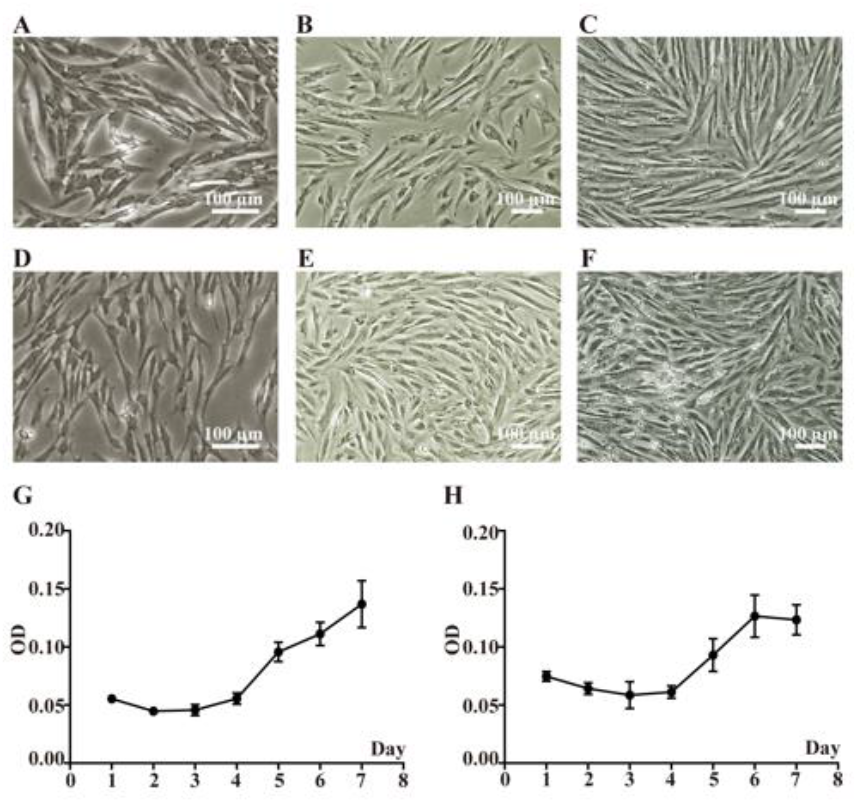
Cultural properties of the mesenchymal stem cell lines. A-C. Representative images of Umbilical Cord-Derived Mesenchymal Stem Cells (UC-MSCs) of primary culture, passage 11 and passage 18 are shown. The cultured cells exhibited a long-thin, spindle-like shape. D-F. Representative pictures of Adipose Tissue-Derived Mesenchymal Stem Cells (AT-MSCs) of primary culture, passage 9 and passage 17 are displayed. The cells also displayed a fibroblastoid morphology and no significant signs of senescence such as cell body enlargement and shape flattening were present. G. Growth curve of UC-MSCs of passage 5 is illustrated. Data are plotted as Mean ± SD. H. Growth curve of AT-MSCs of passage 3 is presented. Data are plotted as Mean ± SD.

In order to measure the proliferation capabilities and morphology change, UC-MSCs and AT-MSCs were subcultured continuously *in vitro*. At passages of 11 and 18, UC-MSCs displayed a bipolar or multipolar shape alike to the primary culture (Figures 1B-1C). Apparent signs of cellular senescence such as enlarged cell bodies and vacuoles were not observed (Figures 1B-1C). Likewise, AT-MSCs could be passaged serially for over 4 months without the occurrence of senescence. AT-MSCs of passages 9 and 17 were of fast propagation and there was no significant change of morphological characteristics (Figures 1E-1F).

In addition, the growth dynamics of the cell cultures were determined based on the plotted growth curves. Although the growth curves did not exhibit a perfect “S”-like shape, they showed an adaptative phase and exponential growth phase (Figures 1G-1H). At the end of the experiment, UC-MSCs seemed to continue proliferating, whilst AT-MSCs reached a plateau phase as anticipated (Figures 1G-1H).

Microorganism contamination could cause a disastrous effect on the cell growth and thus lead to the failure of the cell culture. So as to avoid such unwanted outcomes, high concentrations of antibiotics were added to the solutions to preserve and wash the original tissues. Moreover, the culture medium was supplemented with P/S and amphotericin B during the whole culture period. It turns out that bacterial or fungi growth was not present in the cell cultures based on the direct observations under the light microscope and detections using a BacT/ALERT 3D Microbial Identification System. No cloned DNA bands of mycoplasma were found after nested PCR, indicating the cell cultures were free of mycoplasma contamination.

### 3.2. Chromosome analysis of the cultured MSCs

We collected the UC-MSCs and AT-MSCs for Giemsa staining and the chromosome spreads were analyzed. It is observable that most spreads were composed of 36 chromosomes in total. Because the common hippo is diploid, the chromosomes were paired together according to the G-bands patterns. In details, there were 17 pairs of autosomes and 1 pair of X chromosomes for the cell cultures (Figures 2A-2B). This result is consistent to that of a previous report (Arrighi, 2006). MSCs derived from the umbilical cord and adipose tissue exhibited quite similar banding patterns (Figures 2A-2B). Dramatic chromosome abnormalities such as deletion of large fragments and duplication of a certain chromosome were not found, despite the fact that one or two chromosomes were missing for a few spreads.

**Figure 2.**
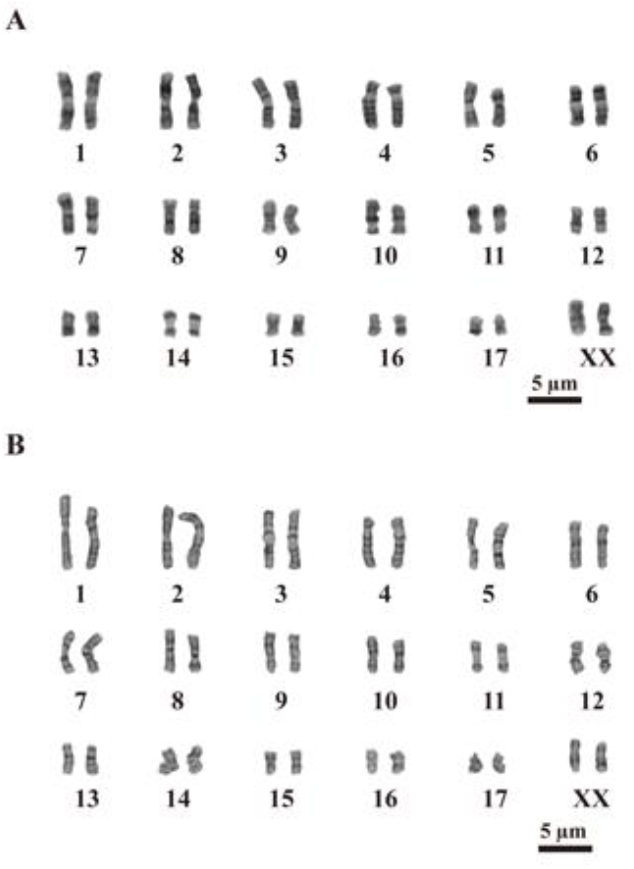
Chromosome analysis of the mesenchymal stem cells. A-B: Karyotyping of UC-MSCs of passage 6 and AT-MSCs of passage 4 is shown. Both cell cultures displayed normal chromosome composition, say, 17 pairs of autosomes and 1 pair of sex chromosomes. No dramatic structural changes were observed.

### 3.3. Expression analysis of the cell surface markers and stem cell related genes

One of the prominent characteristics of MSCs is that they are positive for a few cell surface markers including CD73, CD90, and CD105, but negative for others such as CD45 and HLA-DR. We performed immunofluorescence staining to identify the surface marker expression patterns of our cell lines. Indeed, both UC-MSCs and AT-MSCs expressed CD73, CD90 and CD105 although the positive signals were not as strong as expected (Figures 3A-3B). On the other hand, HLA-DR expression was barely detected (Figures 3A-3B). Nevertheless, these two cell cultures behaved a little differently in terms of the protein expression levels. It seems that the CD73 expression level was higher for UC-MSCs when compared to that for AT-MSCs under the same conditions (Figures 3A-3B). What is most surprising is that both cell cultures were positive for CD45 which is usually considered a leukocyte marker. We speculated that it might be due to the unpredicted influence of a couple of factors on the cell marker expression.

**Figure 3.**
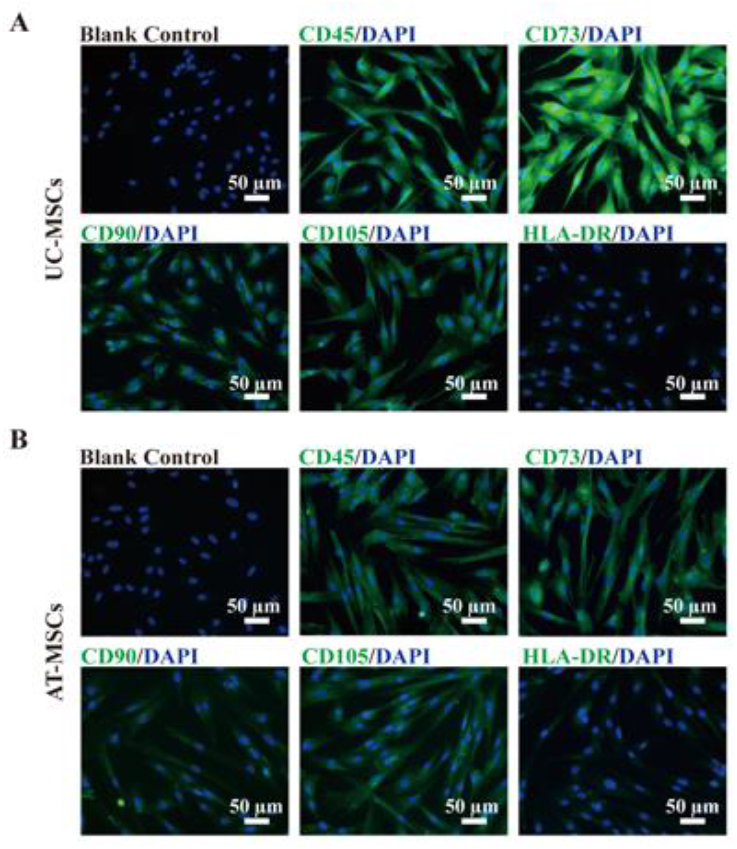
Surface marker expression of the mesenchymal stem cell cultures. A-B: Cell surface markers of UC-MSCs of passage 5 and AT-MSCs of passage 3 were detected with regard to their expression profiles. Both cell lines were strongly positive for CD73, and CD105 and negative for HLA-DR. And they expressed a relatively low level of CD90. It was unexpected that the cell cultures were positive for CD45, a common antigen of hematopoietic cells.

Previous publications indicated that the MSCs were able to express stem cell related proteins (e.g., Deng et al., 2018; Reich et al., 2012; Wei et al., 2016) and thus we examined whether our established cell lines had the same property. In short, UC-MSCs and AT-MSCs were positive for C-MYC, OCT-4 and SOX2 and AT-MSCs seemingly had a slightly higher expression level of these markers (Figures 4A-4B). The expression levels of KLF-4 and NANOG for the cell cultures were pretty low (Figures 4A-4B). We also identified the expression of NESTIN and yet neither of the cultures was detectably positive for this marker (data not shown).

**Figure 4.**
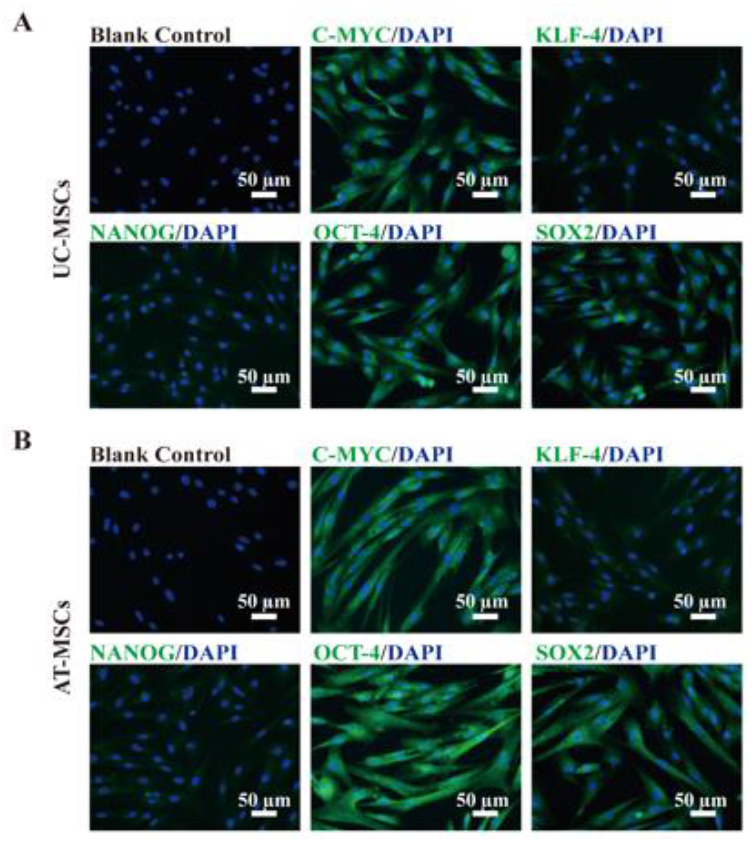
Stem cell related genes expression analysis of the two mesenchymal stem cell lines. A-B. Classical stem cell associated genes expression of UC-MSCs of passage 5 and AT-MSCs of passage 3 was recognized using immunofluorescence staining. The two cell lines were strongly positive for C-MYC, OCT-4 and SOX2, but the expression of KLF-4 and NANOG was barely detected.

### 3.4. Trilineage differentiation of the cultured MSCs

Potential of trilineage differentiation is another essential trait of MSCs. Therefore, we determined the differentiation capabilities of our MSC cultures into three cell types, osteoblasts, adipocytes and chondroblasts. We found that UC-MSCs and AT-MSCs exhibited differential abilities of differentiation under the same induction conditions. There is no doubt that both MSC lines could be induced into chondrocytes because of the positive staining of Alcian blue suggesting the production of glycosaminoglycan component (Figures 5B and 5E). However, when it comes to osteogenic and adipogenic differentiation, the two cell lines displayed contrary results. Positive staining of the UC-MSCs induced cells with Alizarin red indicates the occurrence of osteocytes and thus the differentiation ability of the UC-MSCs (Figure 5A). However, the weak positive staining of Oil Red O suggests the UC-MSCs might not be capable of effectively differentiating into adipogenic lineage in such a circumstance (Figure 5C). On the other side, AT-MSCs had the potential to be induced into adipocytes rather than osteocytes (Figures 5D and 5F).

**Figure 5.**
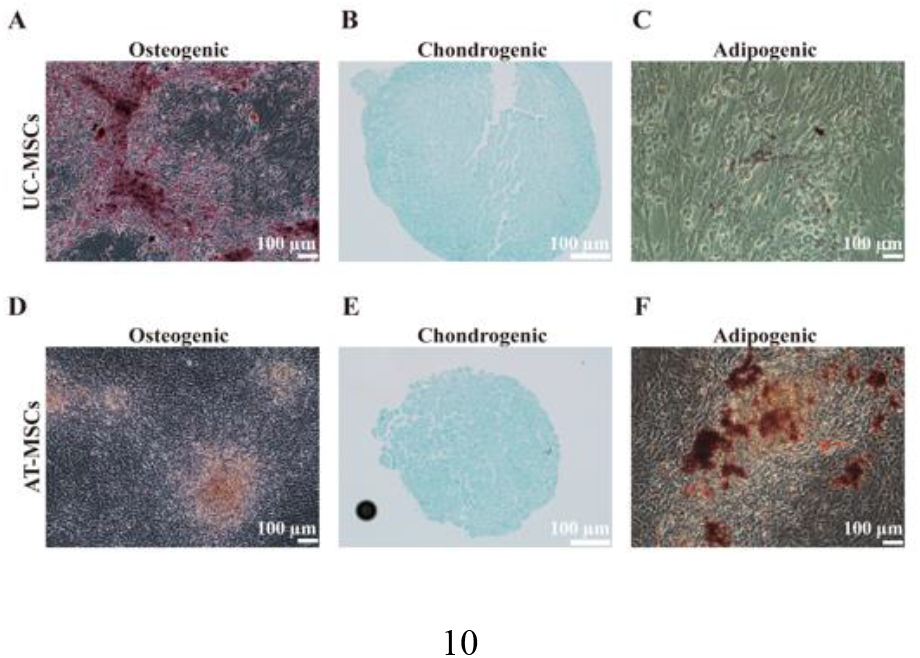
Trilineage differentiation of the mesenchymal stem cells. A-C. Differentiation abilities of the isolated UC-MSCs of passage 5 were identified under the specified induction conditions. They were capable of differentiating into osteocytes and chondrocytes but failed to differentiate into adipocytes. D-F. Differentiation of AT-MSCs of passage 3 was performed under the same conditions as UC-MSCs. AT-MSCs could be induced to adipocytes and chondrocytes and yet osteogenic differentiation could not be generated from AT-MSCs.

## 4. Discussion

MSCs are a population of adult stem cells and present in many tissues of the body. As a result of their therapeutic effects on various diseases, MSCs have been a research hotspot for the last decades. Previous reports showed that MSCs derived from different origins had some common characteristics. Meanwhile, they exhibited heterogeneity regarding the animal species, origin of isolation and culture conditions (Galland and Stamenkovic, 2020; Uder et al., 2018). Expression levels of cell surface markers and stemness associated genes might vary according to the specialized situations (Carrade and Borjesson, 2013). Therefore, identifying the characteristics of MSCs derived from varying animals is necessary and could be helpful for improving the culture system and perceiving their values for future research and applications.

In this project, we established and characterized two new MSC lines derived from the umbilical cord and adipose tissue of a newborn common hippo for the first time to our knowledge. UC-MSCs and AT-MSCs exhibited a fibroblastoid shape and adherent property as what was reported for other MSC cultures. As a large number of MSCs were required for clinical research, *in vitro* expansion of the isolated MSCs was indispensable (Murata et al., 2020). In such case, the proliferation potential of MSCs became a critical factor for their applications. Hence, we serially passaged the UC-MSCs and AT-MSCs to determine their proliferative capacities. Till the end of this study, UC-MSCs and AT-MSCs could be subcultured for 19 passages and 17 passages, respectively. We did not discover significant change of the cellular morphology and their proliferation rate. These results indicated that it is achievable to obtain enough quantity of the cells for downstream research. What is noteworthy is that the population doubling time and colony formation ability might shift when the number of passaging was increased (Deng et al., 2018; Uder et al., 2018). In this sense, it might be needed to measure the proliferation capacity of the cell lines of higher passages.

Genetic stability is another important issue that should be addressed when it is involved in clinical use and animal breeding. As karyotyping is one conventional and effective method to evaluate the possible chromosomal aberrations (Martin and Warburton, 2015; Neri, 2019), we performed chromosome analysis based on G-banding to detect whether there was chromosome abnormality of the cultured MSCs. UC-MSCs and AT-MSCs showed 36 chromosomes in all which were arranged into 18 pairs of them. This result is coincident with the previous discovery (Arrighi, 2006). We did not find abnormal chromosomal alterations in terms of their shapes indicating no large change of the DNA fragments in the order of megabases. It should be noted that karyotyping has a lower resolution when comparing it to other more sensitive techniques such as fluorescent in situ hybridization, single nucleotide polymorphism array and genome sequencing (Martin and Warburton, 2015; Neri, 2019). Karyotyping combined with some other assays may be adopted to discover small molecular defects when these genetic changes are relevant to the therapeutic effects or breeding efficiency.

MSCs are characterized by expression of a few cell surface markers and lack of others. Immunofluorescence staining was applied to test the protein expression panel of the UC-MSCs and AT-MSCs. Expectedly, the cell lines expressed CD73, CD90 and CD105, though the expression level of CD90 was pretty low. Three potential reasons might account for the findings. At first, the antibodies used in the experiments were originally against human cells not common hippo cells and the low expression might be caused by low reactivity between the antibodies and expressed proteins. The second reason could be that the isolated MSCs actually expressed a lower amount of the target proteins when compared to other reported MSCs. This speculation was supported by the research conducted by Wright and his colleagues who found using a species specific antibody against CD90 could not frankly improved the expression level (Wright et al., 2020). Moreover, the culture conditions could have an influence on the immunotype. It was to our surprise that both cell cultures expressed a high level of CD45 while most publications reported a negative reaction. However, a few researchers had the same confusion with us, because their cultured MSCs were also positive for CD45 (Kaiser et al., 2007; Mumaw et al., 2015; Okolicsanyi et al., 2015). We suppose the methods we used for isolation and culture somehow affected the surface marker expression.

At last, induction of UC-MSCs and AT-MSCs into other cell lineages was performed to further detect their stemness maintenance. Both cultures successfully differentiated into chondrocytes as predicted. Nevertheless, UC-MSCs could be induced into osteocytes but not adipocytes and AT-MSCs had opposite abilities. As MSCs are not composed of only one single cell type, they could function heterogeneously and thus had varying differentiation potentials (Galland and Stamenkovic, 2020). Wright et al. also found the low efficiency to differentiate canine UC‐MSCs to adipocytes and they supposed further optimization was likely achievable (Wright et al., 2020). In our case, improving the differentiation protocols and modifying the induction medium might lead to a higher level of differentiation. Differentiation capacities to more cell lineages such as neural cells and germ cells can be another target to be detected in the future.

## 5. Conclusions

In summary, we cultured UC-MSCs and AT-MSCs from a neonatal common hippo and identified their characteristics comprehensively. These two cell lines exhibited major features of MSCs regarding their morphology, expression profiles of the cell surface markers and stem cell related genes and trilineage differentiation abilities. Besides, they had a high cell viability and could be continuously passaged for a couple of months. Their chromosome composition coincides with a previous report of common hippos.

We think the cell cultures could be valuable genetic resources for the downstream research.

## 6. Funding

This research did not receive any specific grant from funding agencies in the public, commercial, or not-for-profit sectors.

## 7. Acknowledgements

We thank Mr. Dongmin Zheng and Mr. Zelong Li for their technical advice and assistance. We are grateful for China National GeneBank DataBase group for their help with the online register of the cell line information.

## 8. Declarations of interest

None.

## 9. Research data availability statement

The data that support the findings of this study have been deposited into CNGB Sequence Archive (CNSA) (https://db.cngb.org/cnsa/faq/) (Guo et al., 2020) of China National GeneBank DataBase (CNGBdb) (Chen et al., 2020) with accession number CNP0002451.

## Notes

### Competing Interest Statement

The authors have declared no competing interest.

